# Precise mapping of new group I introns in tRNA genes

**DOI:** 10.1101/2020.01.27.920355

**Authors:** Kelly P. Williams

## Abstract

Bacterial tRNA have been found interrupted at various positions in the anticodon loop by group I introns, in four types. The primary bioinformatic tool for group I intron discovery is a covariance model that can identify conserved features in the catalytic core and can sometimes identify the typical uridine residue at the -1 position, preceding the 5-prime splice site, but cannot identify the typical guanidine residue at the omega position, preceding the 3-prime splice site, to achieve precise mapping. One approach to complete the automation of group I intron mapping is to focus instead on the exons, which is enabled by the regularity of tRNAs. We develop a software module, within a larger package (tFind) aimed at mapping bacterial tRNA and tmRNA genes precisely, that expands this list of four known classes of intron-interrupted tRNAs to 21 cases. A new covariance model for these introns is presented. The wobble base pair formed by the -1 uridine is considered a determinant of the 5-prime splice site, yet one reasonably large new type bears a cytidine nucleotide at that position.

## INTRODUCTION

Group I introns are self-splicing ribozymes, found interrupting a variety of precursor RNAs. In the first step of splicing, a free guanosine mononucleotide attacks the 5’ splice site, which is typically preceded by a U residue (U^-1^) that forms a wobble base pair in the intron stem-loop P1. In the second step, the G residue (G^Ω^) typically found preceding the 3’ splice site fills the same ribozyme active site as did the free guanosine in step one. Thus the U^-1^ and G^Ω^ nucleotides, if they can be identified in the precursor RNA, precisely delimit the intron. When annotating genomic sequences, the profile Intron_gpI is available from Rfam which locate the catalytic core of the intron and may find the P1 stem-loop at its 5’ end, but does not explicitly search for omegaG. As an alternative to identifying both termini by positive search for the intron, it may be possible to precisely map its two surrounding exons, especially when the mature RNA is highly regular, as are tRNAs. Some number of group I introns are known to reside in the anticodon loop of bacterial pre-tRNAs (1-3). The tRNA-like domain of the bacterial tmRNA, which lacks an anticodon loop, is occasionally interrupted by a group I intron, in its T-loop. Here we apply tRNA/tmRNA annotation software and the Rfam group I intron profile to discover and precisely map many new group I introns in bacterial tRNA genes. We then build a new profile, which in a second iteration expands the discovery further.

## MATERIALS AND METHODS

The nucleotide sequence dataset used was the 80413 bacterial and 1028 archaeal genome assemblies available from GenBank in January 2017 that had ≤ 300 scaffolds and N50 ≥ 10000. The covariance model (CM) used in discovery phase I (see text) was Intron_gpI from Rfam (4). To build a new CM for phase II, 26 representative bacterial introns were selected interrupting tmRNAs and the tRNAs found in phase I, choosing shorter introns less likely to contain extraneous loop insertions. Sequences began with the 5’ P1 stem that contains U^-1^ and continued to the G^Ω^, beyond which little conservation was apparent. Alignment was primed using *cmalign* from the Infernal 1.1.2 package (5), and followed by manual alignment to best fit the conserved secondary structure features (Supp. File gpI.sto). The new CM was built using *cmbuild* from Infernal (Supp. File gpI_bact_tRNA.cm).

## RESULTS AND DISCUSSION

### tFind software

Careful mapping and annotation of tRNA and tmRNA genes was motivated by our programs for finding genomic islands in such genes (6) (manuscript submitted). Our package tFind searches for tmRNA genes and for tRNA genes interrupted or not by group I introns (Fig. 1). Three scripts apply different approaches to finding tmRNA genes: i) the program Aragorn (7), ii) application of four tmRNA covariance models (CMs) from Rfam (4), and iii) a BLAST-based approach using a large curated tmRNA database. This latter script, rfind.pl, uses iterative genome masking, allowing discovery of secondary tmRNA genes whose signals may be much weaker than a primary gene. Together these approaches perform well to identify tmRNA genes of all three major types (standard, two-piece, and group I intron-bearing). Application of tFind to the genome set yielded 343 tmRNA genes with a group I intron interrupting the T stem-loop as described previously (8). Precise tmRNA gene mapping comes from rfind and (for standard tmRNAs) from Aragorn. Although only genes matching perfectly to the curated database advance to the final document (ttm.gff), the system is meant to be extensible, such that convincing novel tmRNA genes discovered among the rejects.gff outputs or by other means can be added to the database for improved future performance. tRNA gene-finding relies on the well-known tRNAscan-SE (1), whose latest version applies covariance models for each tRNA isotype family, separately for Archaea and Bacteria; we add careful demarcation of the discriminator position and inclusion of the tRNA-His G^-1^ nucleotide when present. We detail below the function and yield from the introns.pl component that finds tRNA genes interrupted by group I introns. A final pass resolves overlaps arising from these components. The tFind package is available at github.com/sandialabs/TIGER.

**Fig. 1.**
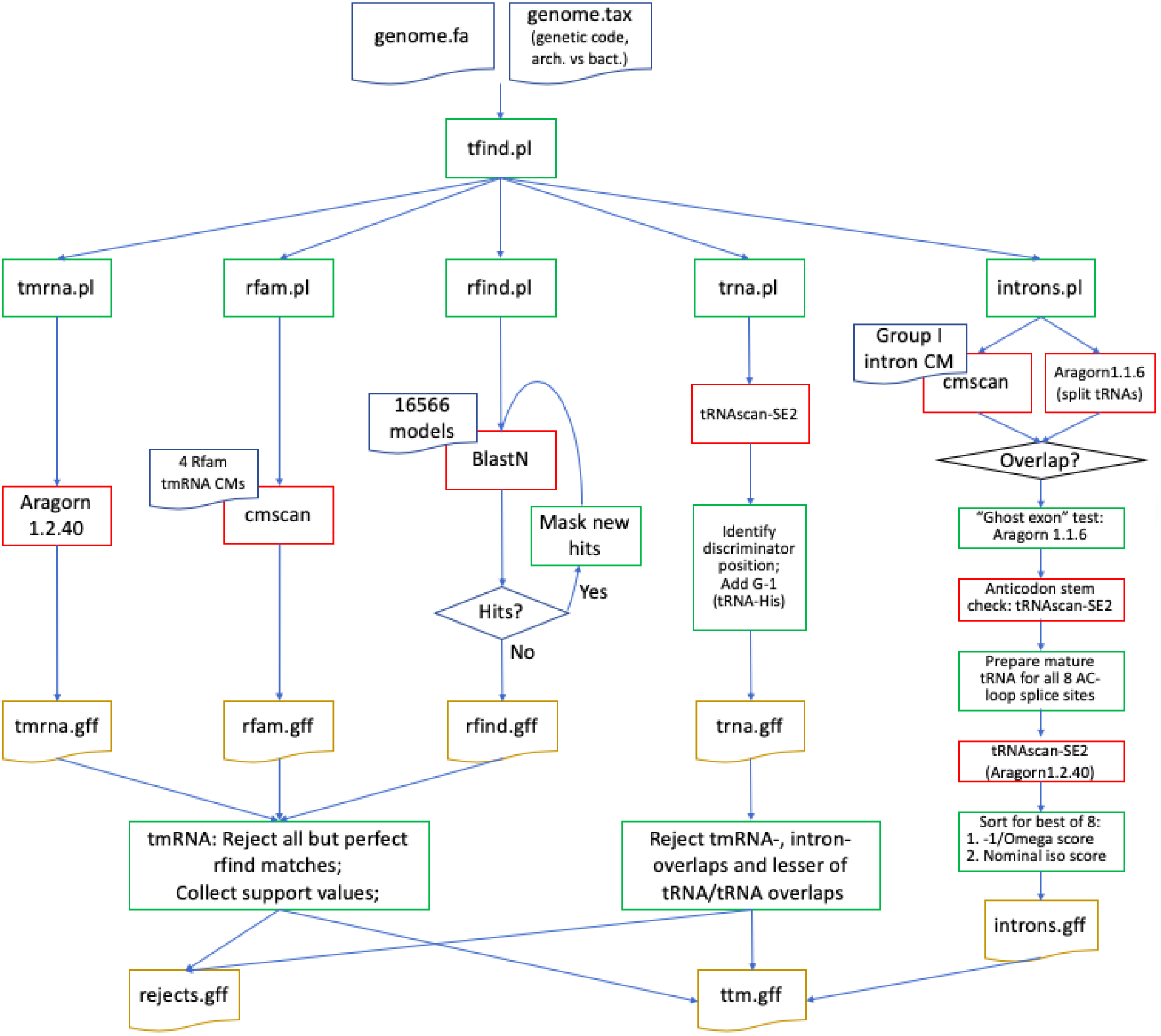
tFind pipeline for precise mapping of tRNA and tmRNA genes in bacterial genomes. Three avenues tackle tmRNA genes, one is for uninterrupted tRNA genes and introns.pl identifies tRNA genes interrupted by group I introns.

### Intron-mapping software

The pipeline for precise mapping of group I introns within tRNA genes is shown at the right of Fig. 1. The Rfam CM Intron_gpI is applied to the genome, and if there are any above-threshold (20 bits) hits, Aragorn 1.1.6 is run so as to allow detection of an intron of up to 3 kbp within the anticodon loop. Cases where a group I intron call maps within a split tRNA call are processed further. Eight possible intron insertion sites are considered, on either side of each of the 7 anticodon loop nucleotides, and the corresponding eight hypothetical mature RNA sequences are generated. These are scored three ways, with: i) a group I intron splice site “-1/ Ω” score where the 5’-preceding nucleotide adds 2 to the score if a U or 1 to the score if a C (based on our Phase I observation of its use in convincing introns), and the Ω (3’-terminal) position of the candidate intron adds 3 to the score if a G, ii) the isotype score according to tRNAscan-SE, and iii) the “IPD” score from tRNAscan-SE reflecting mismatch between the nominal amino acid identity based on the anticodon nucleotides and the isotype profile CM match. The splice site is selected among the eight by sorting by each of the three scores in the given order of precedence.

We found that Aragorn would occasionally call false “ghost” exons upstream or downstream of the actual ones, and could usually solve this problem by testing the primary call again after deleting the called 5’ exon, 3’ exon or both, and proceeding with the best scoring of the four. We noticed that certain introns that had been miscalled because the wrong 3’ partner sequence for the anticodon stem had been chosen by Aragorn. We therefore gave tRNAscan-SE an opportunity to correct the anticodon stem call, highly truncating the intron so that tRNAscan-SE could process it, with only 12 ntd from each end of the intron. For a small number of cases tRNAscan-SE did not recognize the spliced tRNA, so a backup subroutine employing Aragorn 1.2.40 was developed for such cases.

Regarding nomenclature, we number the 8 tested anticodon positions 0-7, 5’ to 3’, and note the-1 base and the Ω base, preceding with a unique identifier for the genome assembly; thus the classical intron invading tRNA-Leu in *Nostoc azollae* (3), spliced out between the slashes in CUU/a…g/AAAA (mature anticodon loop sequence in uppercase) is Naz2.Leu.3Ug.

### Phase I discovery

Introns.pl was run with the Rfam CM Intron_gpI on 80413 bacterial and 1028 archaeal genomes. This yielded 10174 hits to Intron_gpI scoring ≥ 18 bits (well below its suggested “gathering cutoff” of 40 bits), that came from 6734 genomes. For these genomes Aragorn 1.1.6 called 11016 tRNA genes with introns ≥ 50 bp in the anticodon loop. There were 4406 cases of Intron_gpI calls within split tRNA gene calls, from 3879 genomes, producing candidate gpI intron-split tRNA genes. All the genomes involved were bacterial, except one named “Candidatus *Pacearchaeota* archaeon RBG_16_35_8”. This genome had been prepared by binning scaffolds from a metagenomic assembly (9). Rather than suggesting that we have discovered the first known group I intron for an archaeon, we suspect that this tRNA gene was from a bacterial scaffold mis-binned with archaeal scaffolds in that project. Indeed 14 of the 21 raw hits to Intron_gpI for nominally archaeal genomes came from that same project.

### Phase II discovery

None of the 12 seed sequences used to build the original Rfam covariance model Intron_gpI were bacterial. A new CM gpI_bact_tRNA was prepared, seeded only with introns from bacterial tRNA or tmRNA genes, extending from U^-1^ to G^Ω^. Its use in a new run of introns.pl improved the yield by 20.2%, to 5292 introns, missing none of those found in phase I.

Table 1 shows the breakdowns in yields by tRNA interruption site and bacterial phylum. Cyanobacteria have the highest frequency, 0.69 interrupted tRNA genes per genome, followed by the metagenomic assemblies from unnamed phyla, with 0.49 per genome.

**Table 1.** Distribution by phylum and by tRNA isoacceptor/anticodon loop setting, for the 5292 split tRNA genes found in phase II and the 4406 found in phase I. Values are for the phase II results using the new CM, except for the column and row marked “Rfam CM” reporting phase I counts. Blue shading marks the four classical cases of interrupted bacterial tRNA genes and green marks 17 new ones; yellow marks erroneous calls with suggested corrections given. Settings that are not U^-1^/G^Ω^ are shown in red type.

| Intron setting | Phylum |  |  |  |  |  |  |  |  |  |  |  |  |  |  |  |  | Correct to | Notes |
| --- | --- | --- | --- | --- | --- | --- | --- | --- | --- | --- | --- | --- | --- | --- | --- | --- | --- | --- | --- |
|  | Unamed | Acidobacteria | Actinobacteria | Bacteroidetes | Chlorobi | Chloroflexi | Cyanobacteria | Elusimicrobia | Fibrobacteres | Firmicutes | Nitrospirae | Planctomycetes | Proteobacteria | Verrucomicrobia | 21 other phyla | All Bacteria | All (Rfam CM) |  |  |
| Leu.3Ug | 3 | 9 |  | 11 |  | 2 | 142 |  |  | 8 |  | 2 | 2386 |  |  | 2563 | 2393 |  |  |
| Asn.4Ug | 554 | 1 |  |  |  | 2 | 6 | 12 |  | 6 | 3 | 1 | 32 |  |  | 617 | 586 |  |  |
| Arg.5Ug |  |  | 1 |  |  | 1 | 85 |  |  |  |  |  | 475 |  |  | 562 | 460 |  |  |
| Ser.3Ug |  |  |  |  |  |  |  |  |  |  | 1 |  | 212 |  |  | 213 | 5 |  |  |
| fMet.2Ug | 133 |  |  | 3 | 1 |  | 35 | 4 |  |  | 4 |  | 23 |  |  | 203 | 199 |  |  |
| Arg.3Ug | 3 | 4 |  | 7 |  |  | 1 | 2 |  | 126 |  |  | 44 |  |  | 187 | 55 |  |  |
| Thr.5Ug | 106 |  |  |  |  |  |  |  |  | 6 |  |  | 58 |  |  | 170 | 61 |  |  |
| Ile2.2Ug | 41 | 1 |  |  |  | 1 |  | 1 |  | 1 |  | 1 | 121 | 2 |  | 169 | 125 |  |  |
| Phe.2Ug | 128 |  |  |  |  |  |  |  |  |  |  |  |  |  |  | 128 | 126 |  |  |
| Ile2.5Ug | 6 |  |  | 4 |  |  |  |  |  |  |  | 7 | 108 |  |  | 125 | 121 |  |  |
| Asp.4Ug | 73 |  |  |  |  |  |  |  |  |  |  |  |  |  |  | 73 | 65 |  |  |
| Leu.3Cg |  |  |  |  |  |  |  |  |  |  |  |  | 72 |  |  | 72 | 43 |  |  |
| fMet.5Ug | 54 |  |  |  |  |  |  |  |  |  |  |  |  |  |  | 54 | 53 |  |  |
| Met.2Ug | 37 |  |  |  |  |  |  |  |  |  |  |  |  |  |  | 37 | 35 |  |  |
| Lys.3Ug |  |  | 25 |  |  |  |  |  |  |  |  | 7 |  |  |  | 32 | 22 |  |  |
| Tyr.4Ug | 17 |  |  |  |  |  |  |  |  | 9 |  |  |  |  |  | 26 | 14 |  |  |
| Lys.2Ug | 2 |  |  |  |  |  |  |  |  |  |  |  | 8 |  |  | 10 | 10 |  |  |
| Lys.4Ug | 2 | 1 |  |  |  |  |  |  |  |  |  |  | 5 |  |  | 8 | 5 |  |  |
| Thr.2Ug | 8 |  |  |  |  |  |  |  |  |  |  |  |  |  |  | 8 | 7 | Asn.4Ug | Isoacceptor score slightly better for incorrect 2Ug |
| Thr.3Ug |  |  |  |  |  |  |  |  |  |  |  |  | 8 |  |  | 8 | 8 |  |  |
| Sup.3Ug |  |  |  |  |  |  |  |  | 7 |  |  |  |  |  |  | 7 | 0 | Ile2.5Ug | Problematic ac stem-loop: GTcggACTCAT/TAAACccgTT |
| Leu.1Ug |  |  |  |  |  |  |  |  |  |  |  |  | 5 |  |  | 5 | 5 |  |  |
| Met.5Ug | 5 |  |  |  |  |  |  |  |  |  |  |  |  |  |  | 5 | 0 | Ile2.5Ug | Fails tRNAscan-SE; all 5 identical |
| Leu.2Ug |  |  |  | 1 |  |  |  |  |  |  |  |  | 3 |  |  | 4 | 3 |  |  |
| Asn.3Ag | 1 |  |  |  |  |  |  |  |  |  |  |  |  |  |  | 1 | 1 | omit | Genomic deletion of critical tRNA gene segment |
| fMet.2Cg | 1 |  |  |  |  |  |  |  |  |  |  |  |  |  |  | 1 | 1 | fMet.2Ug | 8 nt ac loop |
| Ile2.5Uu |  |  |  |  |  |  |  |  |  |  |  |  | 1 |  |  | 1 | 1 | Ile2.2Ug | Worse score after removing ghost 5' and 3' exons |
| Leu.0Ag |  |  |  |  |  |  |  |  |  |  |  |  | 1 |  |  | 1 | 1 | Leu.1Ug | 8 nt ac loop |
| Undet.3Ug |  |  |  |  |  |  |  |  |  | 1 |  |  |  |  |  | 1 | 0 | Arg.3Ug | Very similar to Arg.3Ug but changes in P1 favor Arg.vLoop |
| Undet.4Un | 1 |  |  |  |  |  |  |  |  |  |  |  |  |  |  | 1 | 1 | Leu.3Ug | Ambiguous bases in anticodon region |
| Genes | 1175 | 16 | 26 | 26 | 1 | 6 | 269 | 19 | 7 | 157 | 8 | 18 | 3562 | 2 | 0 | 5292 |  |  |  |
| Genes(Rfam CM) | 1008 | 11 | 22 | 21 | 1 | 6 | 264 | 19 | 0 | 16 | 7 | 11 | 3020 | 0 | 0 |  | 4406 |  |  |
| Genomes | 2420 | 88 | 10058 | 1920 | 22 | 175 | 388 | 56 | 29 | 27537 | 71 | 80 | 35196 | 81 | 1837 | 79958 |  |  |  |
| Genes/genome | 0.486 | 0.182 | 0.003 | 0.014 | 0.045 | 0.034 | 0.693 | 0.339 | 0.241 | 0.006 | 0.113 | 0.225 | 0.101 | 0.025 | 0 | 0.066 |  |  |  |

Validation comes from a database of group I introns (10), which lists only the four classical varieties in bacterial tRNAs (Leu.3Ug, Arg.5Ug, fMet.2Ug and Ile2.5Ug), and marks them all as subgroup IC3 (11); these four categories totaled 3453 introns, or 65.2% of all discovered introns. The maximum intron length detectable was 3000 bp, yet the longest intron found was 1817 bp and the second longest was 922 bp (Fig. 2). Further independent validation comes from the finding of P1 stems containing U^-1^ in a wobble base pair.

**Fig. 2.**
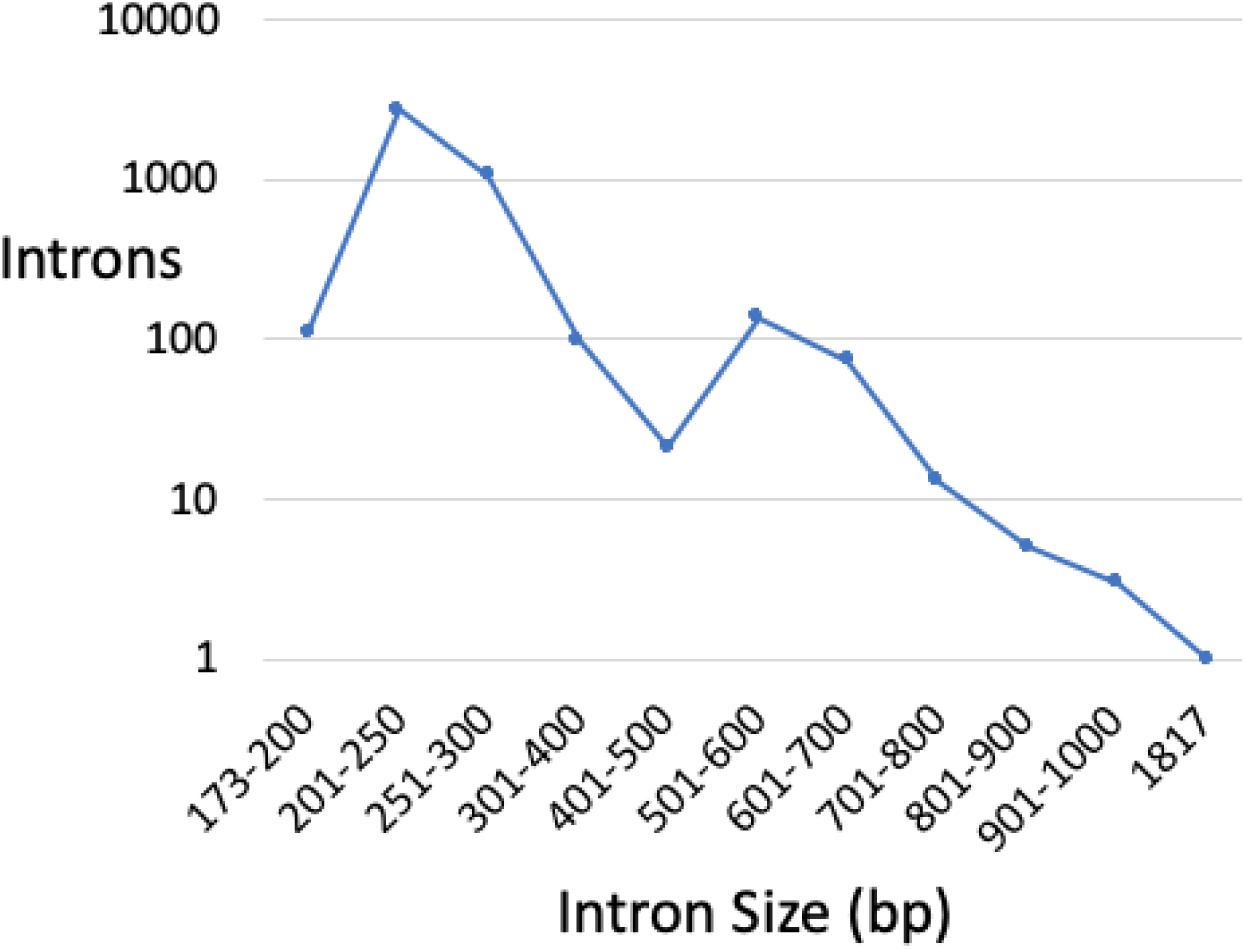
Intron sizes detected. Those above ∼400 bp may contain whole or partial protein-coding genes.

Finally another validation is that a U^-1^/g^Ω^ pair could be found at one of the 8 tested anticodon loop positions in a very high fraction (98.5%) of the raw calls. This value would be unlikely to arise by chance; the expected occurrence of any given nucleotide pair among 8 random pair selections is 1-(15/16)^8^=40.3%. The 76 raw calls with no U^-1^/g^Ω^ pair revealed four cases of various remaining errors for introns.pl or in the genome assembly, but also a relatively large type of 72 Leu.3Cg introns, all found in Proteobacteria, which forms a Watson-Crick C^-1^:G base pair in stem P1 rather than the usual U^-1^:G. Such mutation inactivates the *Tetrahymena* group I intron (12), so the ability of the Leu.3Cg pre-tRNAs to splice needs testing.

## Conclusion

We have added many new cases of tRNA genes split within the anticodon loop region by group I introns (Fig. 3) and provide software that can be applied for such discovery to any new bacterial genomic or metagenomic sequence data. The error rate we measure for false positives and misidentification of the splice site is 26/5296 = 0.50%.

**Fig 3.**
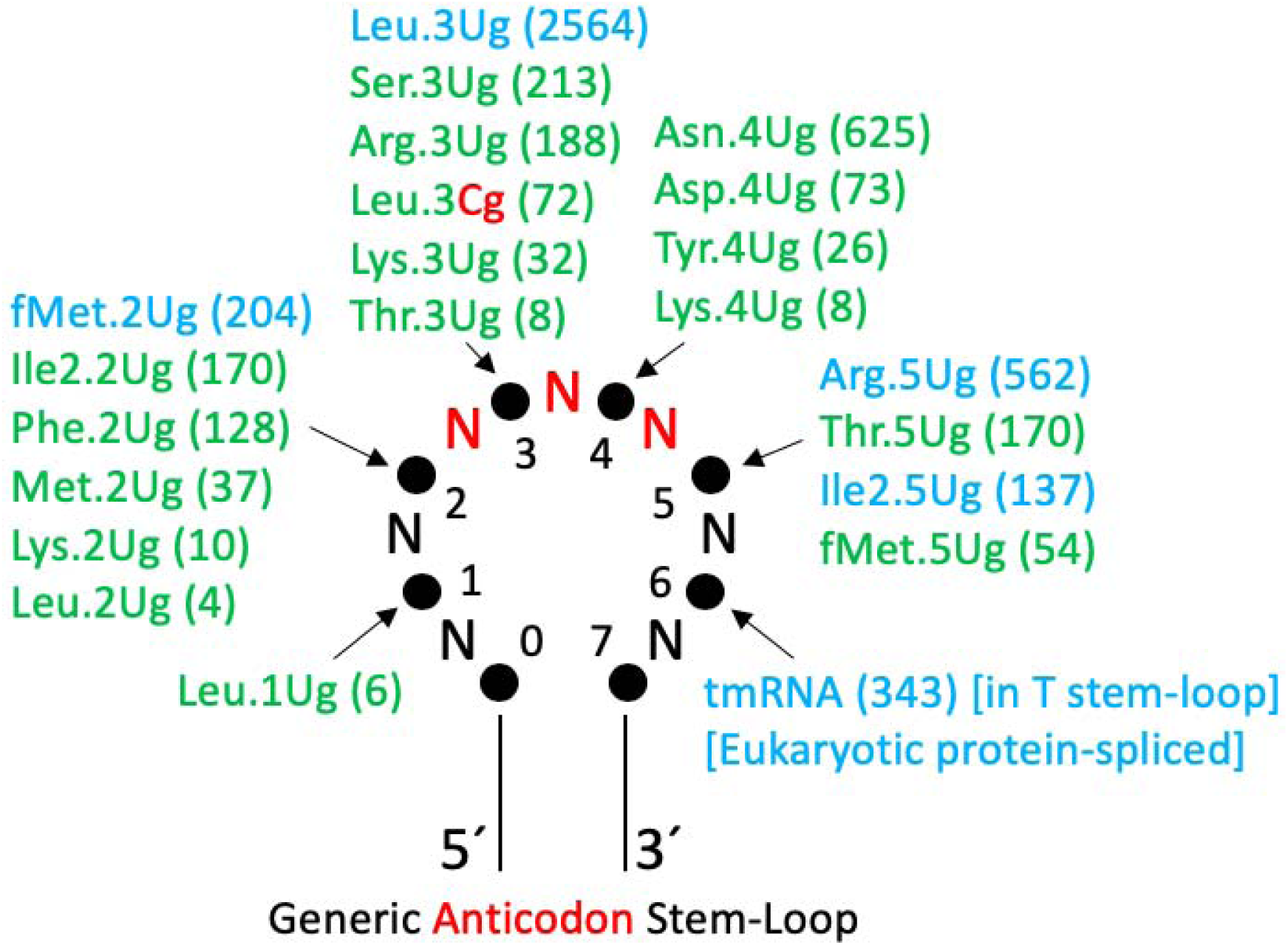
Summary of intron locations. Data (counts in parentheses) from Table 1, with the same coloring scheme are showed here, after making the small number of corrections noted in the Table. The eight tested positions in the anticodon loop are numbered. Also shown are the site of the tmRNA group I introns and eukaryotic protein-spliced introns.

## Supporting information

Supp. File gpI.sto

Supp. File gpI_bact_tRNA.cm

## ACKNOWLEDGEMENTS

We thank Dean Laslett for kind reconfiguration of Aragorn and allowing us to include its code and executables in the package.

